# Holotomography reveals biophysical remodeling of mouse oocytes during post-ovulatory aging

**DOI:** 10.64898/2026.06.18.733271

**Authors:** Minsu Kim, Chungha Lee, Jieun Do, Taeseop Shin, Daria Iliev, Kyoung Hee Choi, So Yeon Shin, Jieun Ko, Teodora Popa, Cristina Hickman, Ji Hyang Kim, YongKeun Park

## Abstract

Routine assessment of oocyte state in reproductive biology and clinical embryology still relies largely on two-dimensional transmitted-light morphology, leaving the underlying three-dimensional intracellular organization largely unmeasured. Here, we use holotomography to reconstruct volumetric refractive index (RI) distributions in unlabeled live mouse oocytes and combine this imaging with sparse-annotation, AI-assisted compartment segmentation and multidimensional feature extraction across geometry, RI statistics, dry-mass density, and intracellular texture. Applied to an *in vitro* model of post-ovulatory aging, holotomography-derived biophysical profiles separated fresh and aged oocytes more strongly than two-dimensional brightfield-derived features, and feature-family ablation indicated that non-morphological descriptors contributed substantially to this separation. At the population level, aged oocytes showed coordinated ooplasmic remodeling, including reduced volume, increased RI heterogeneity, and increased dry-mass density while total dry mass was broadly conserved. These findings demonstrate a label-free quantitative framework for detecting post-ovulatory aging-associated biophysical remodeling in intact mammalian oocytes.

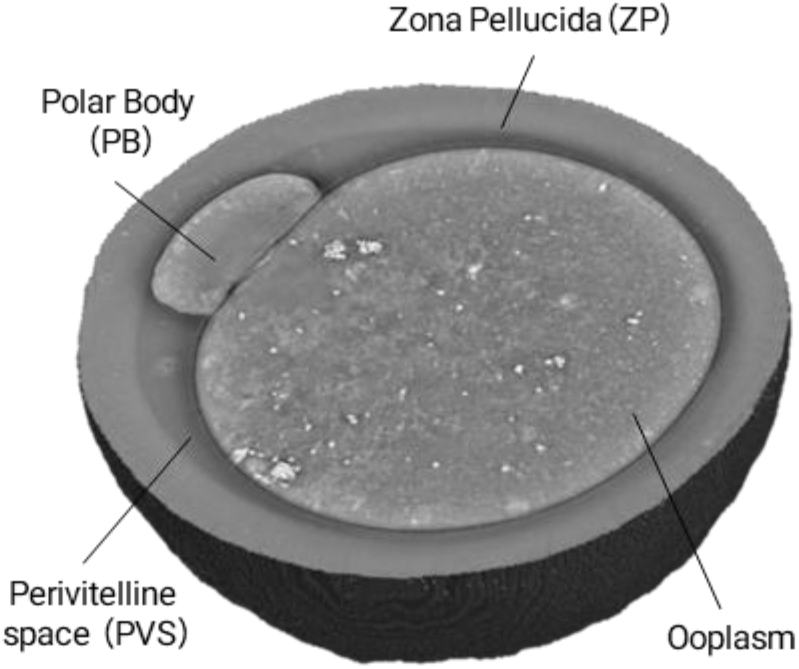

## Introduction

Oocyte quality is a central determinant of fertilization and early embryonic development^1,2^, yet evaluation in oocyte biology research and clinical embryology still relies largely on transmitted-light microscopy for morphological assessment — simple, rapid, and non-destructive, but limited in its ability to resolve internal three-dimensional (3D) organization^3^. Features likely relevant to oocyte state, including 3D intracellular organization, cytoplasmic heterogeneity, and subcellular remodeling, are not readily accessible to routine inspection^4,5^. Computational transmitted-light phenotyping pipelines have shown that engineered features extracted from brightfield (BF) images can classify oocyte populations and maturation potential^5^, but such descriptors remain bounded by the 2D intensity contrast of BF acquisition. A non-invasive, quantitative approach to the internal biophysical organization of intact oocytes would therefore address an important gap in reproductive biology. Here, we ask whether label-free volumetric refractive index (RI) imaging can quantitatively capture the internal state of intact mouse oocytes and what such measurements add beyond BF analysis.

Several label-free modalities have been applied to living mammalian oocytes and early embryos, each accessing a distinct physical contrast. Back-scattering approaches, including full-field and ultrahigh-resolution optical coherence microscopy, optical coherence tomography, and reflection-matrix microscopy, can resolve nuclear and cytoskeletal structures and, in the oviduct, even intravital transport^4,6–9^. Nonlinear and vibrational imaging, including higher-harmonic generation, coherent anti-Stokes Raman scattering (CARS), and spontaneous Raman microscopy, provides structural or chemical contrast through nonlinear susceptibility and molecular vibrations^11–13^. Brillouin microscopy maps mechanical stiffness^10,11^, whereas biodynamic digital holographic speckle microscopy extracts intracellular dynamic-scattering spectra as functional readouts of viability^12,13^. These modalities access complementary axes, back-scattering, nonlinear susceptibility, chemistry, mechanics, or dynamics, but do not directly provide the quantitative volumetric RI distribution from which dry mass, compartment-resolved RI statistics, and 3D intracellular texture can be derived in a single label-free acquisition (Supplementary Table 1).

A complementary axis is provided by label-free RI imaging, where RI, an intrinsic optical property related to local composition and mass density, carries quantitative structural information without exogenous dyes or destructive processing. Holotomography (HT) reconstructs volumetric RI distributions from multiple intensity measurements and resolves intact biological samples in 3D, without labels or photobleaching^14–23^. Despite this promise, it remains unclear whether HT can support systematic quantitative phenotyping of oocytes and whether RI-derived features provide information beyond BF analysis^24^. A framework combining label-free 3D imaging of intact oocytes with compartment-resolved segmentation and multidimensional biophysical feature extraction has not yet been established, and the feature classes that may underlie any advantage over conventional analysis remain unknown.

Here, we build an HT-based phenotyping framework for intact mouse oocytes. Volumetric RI reconstruction resolves the ooplasm, zona pellucida (ZP), perivitelline space (PVS), and polar body (PB), and an artificial intelligence (AI)-assisted segmentation pipeline supports compartment-level feature extraction across morphology, dry mass, RI statistics, and intracellular texture. Applied to an *in vitro* model of post-ovulatory aging^25,26^, the framework captures state-dependent intracellular reorganization and separates aging states more effectively than BF analysis. Ablation-based analysis identifies the feature classes that contribute to this advantage, supporting HT as a quantitative readout of aging-associated biophysical remodeling rather than a generic 3D imaging technique.

## Results

### Workflow overview and label-free 3D imaging of intact mouse oocytes

We developed a four-stage pipeline for label-free multidimensional biophysical phenotyping of intact mouse oocytes, combining HT, AI-assisted compartment segmentation, and RI-derived feature extraction across post-ovulatory aging conditions (**Fig. 1**). Each oocyte was imaged on an HT system under patterned illumination, reconstructed as a volumetric RI tomogram, segmented into four structural compartments (ooplasm, PVS, ZP, and PB) and represented as a multidimensional biophysical profile spanning morphology, dry mass, RI statistics, and intracellular texture for comparison between fresh and in vitro post-ovulatory aged oocytes.

**Fig. 1.**
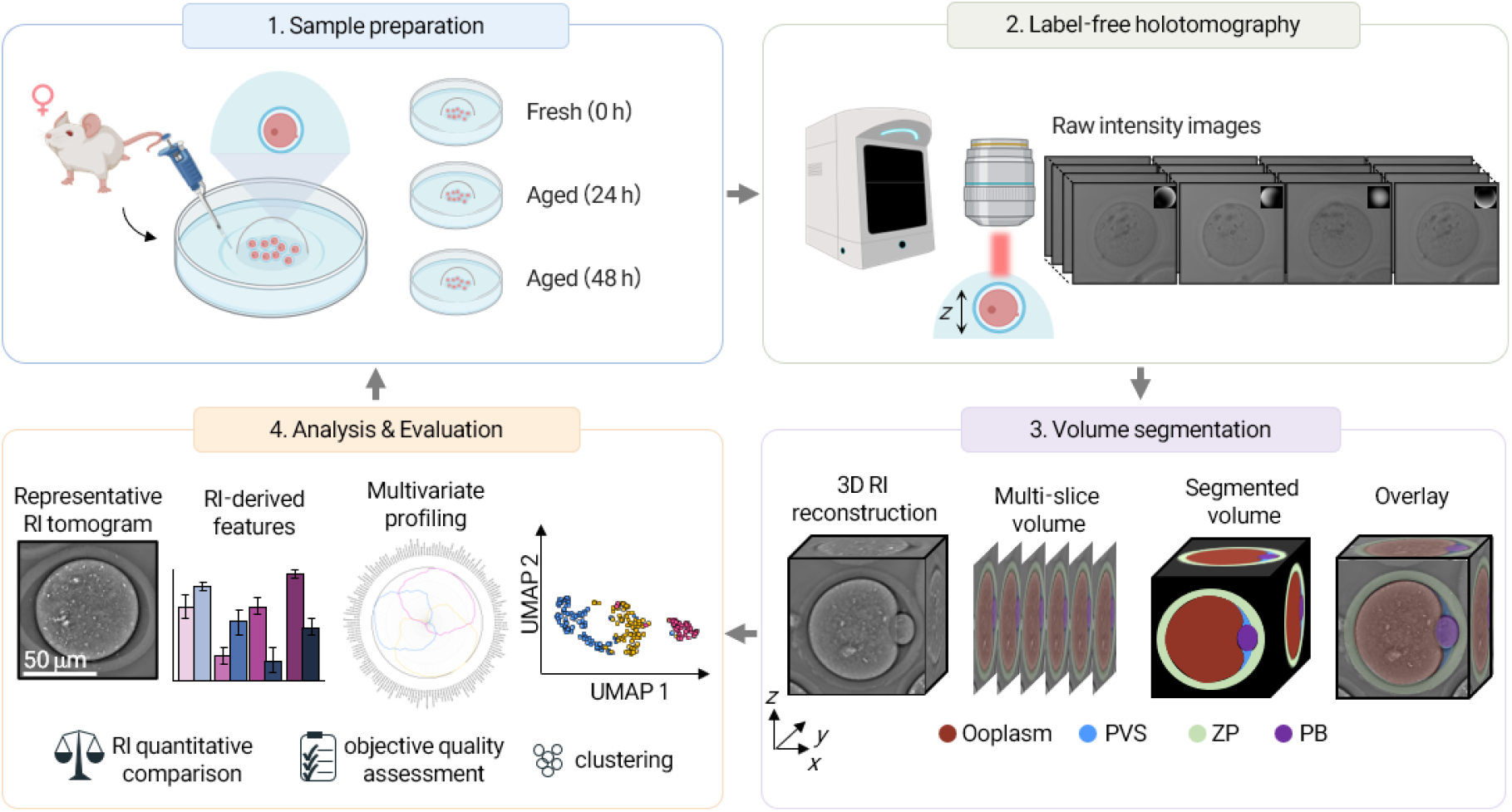
Workflow for label-free multidimensional biophysical phenotyping of mouse oocytes. A four-stage pipeline consists of four steps: sample preparation, label-free holotomography, volume segmentation, and analysis and evaluation. During sample preparation, MII oocytes are retrieved from superovulated female mice and assigned to three groups: fresh (0 h), aged-24 h, or aged-48 h, corresponding to 0, 24, and 48 h of in vitro culture. Each oocyte is then imaged using label-free holotomography under patterned illumination to acquire raw intensity images. For volume segmentation, a 3D RI tomogram is reconstructed from the acquired intensity images, and each oocyte volume is partitioned into four structural compartments, ooplasm, perivitelline space (PVS), zona pellucida (ZP), and polar body (PB), using a SAM 2-based propagation pipeline. In the analysis and evaluation step, compartment-level RI-derived features spanning morphology, dry mass, RI statistics, and intracellular texture are extracted for downstream analysis.

Label-free HT visualized the 3D architecture of intact mouse oocytes with subcellular detail. Volumetric RI reconstructions delineated the PB, ZP, PVS, and ooplasm simultaneously without exogenous labelling (**Fig. 2a**), and axial RI slices through the same oocyte at successive depths revealed RI-dense intracellular structures within the ooplasm as well as fine-scale substructure within the ZP (**Fig. 2b**). HT thus captured both bulk compartmental architecture and depth-resolved subcellular organization in a single label-free acquisition.

**Fig. 2.**
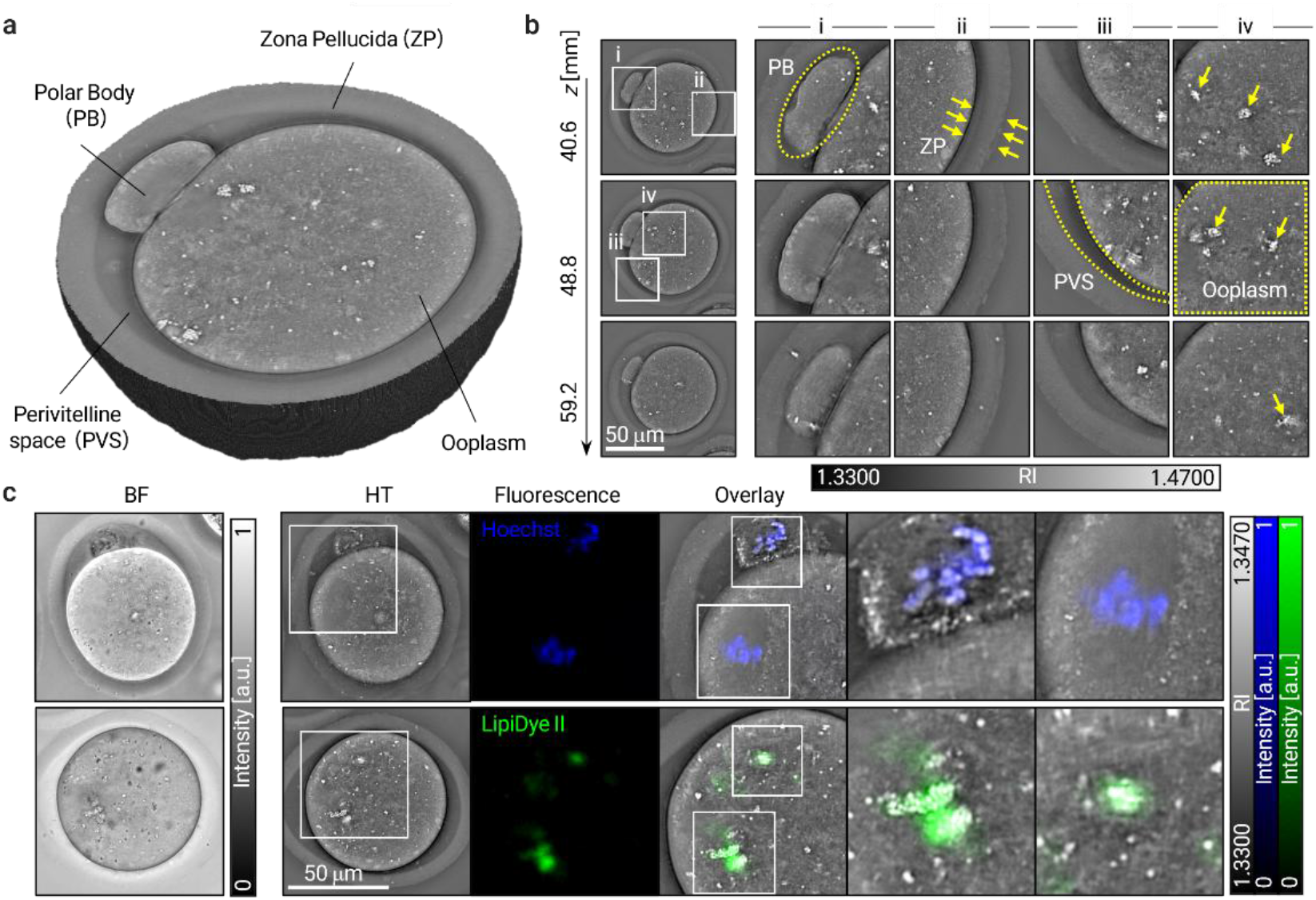
Label-free holotomography reveals 3D architecture and subcellular organization of intact mouse oocytes. **a**, Volumetric RI rendering of a representative MII mouse oocyte. PB, ZP, PVS, and ooplasm are resolved simultaneously in three dimensions. **b**, Axial RI slices through the same oocyte at successive depths. Z positions are indicated on the left and correspond to the slice depth within the reconstructed volume. The first column shows the full in-plane view at each depth; columns 2–5 show zoomed views of the boxed regions in column 1, highlighting representative subcellular structures across compartments. Yellow dashed outlines mark representative compartments; yellow arrows indicate the ZP boundary and RI-dense features within the ZP and ooplasm. **c**, Correspondence between HT contrast and fluorescence labelling in the same oocytes. Top row, Hoechst labels chromatin at the metaphase plate. Bottom row, LipiDye II labels lipid-rich structures. Columns show, from left to right, brightfield (BF), the in-focus HT slice, fluorescence signal, and zoomed HT-fluorescence overlays. White boxes indicate regions shown at higher magnification. Overlays show spatial correspondence between RI-dense intracellular regions and fluorescence signals.

To assess whether this RI contrast reflects biologically meaningful intracellular organization, we compared HT with BF and fluorescence imaging of the same oocytes. HT showed intracellular features with higher contrast than BF, and RI-dense regions overlapped spatially with fluorescence signals from Hoechst, marking chromatin at the metaphase plate, and LipiDye II, marking lipid-rich structures (**Fig. 2c**; HT-Hoechst and HT-LipiDye II overlays). These observations suggest that RI distributions reflect biologically relevant intracellular organization rather than non-specific optical variation, supporting the quantitative compartment-level analysis developed below.

### RI imaging reveals stage-specific and aging-associated patterns in mouse oocytes

HT revealed state-associated intracellular RI organization across oocyte maturation and post-ovulatory aging conditions. Representative oocytes at germinal vesicle breakdown (GVBD), metaphase I (MI), and metaphase II (MII) displayed stage-specific intracellular RI patterns that were not apparent in the corresponding BF images (**Fig. 3a**). Across three independent post-ovulatory aging groups: fresh (0 h), aged-24 h, and aged-48 h, HT images of distinct representative MII oocytes showed condition-associated differences in intracellular RI organization (**Fig. 3b**). In the in vitro post-ovulatory aging model^25,26^, freshly collected oocytes showed more coherent intracellular RI organization, whereas oocytes cultured for 24 h or 48 h showed greater RI texture heterogeneity, RI-dense intracellular regions, and irregular ooplasmic patterns. These condition-associated differences were more apparent in HT than in the corresponding BF images.

**Fig. 3.**
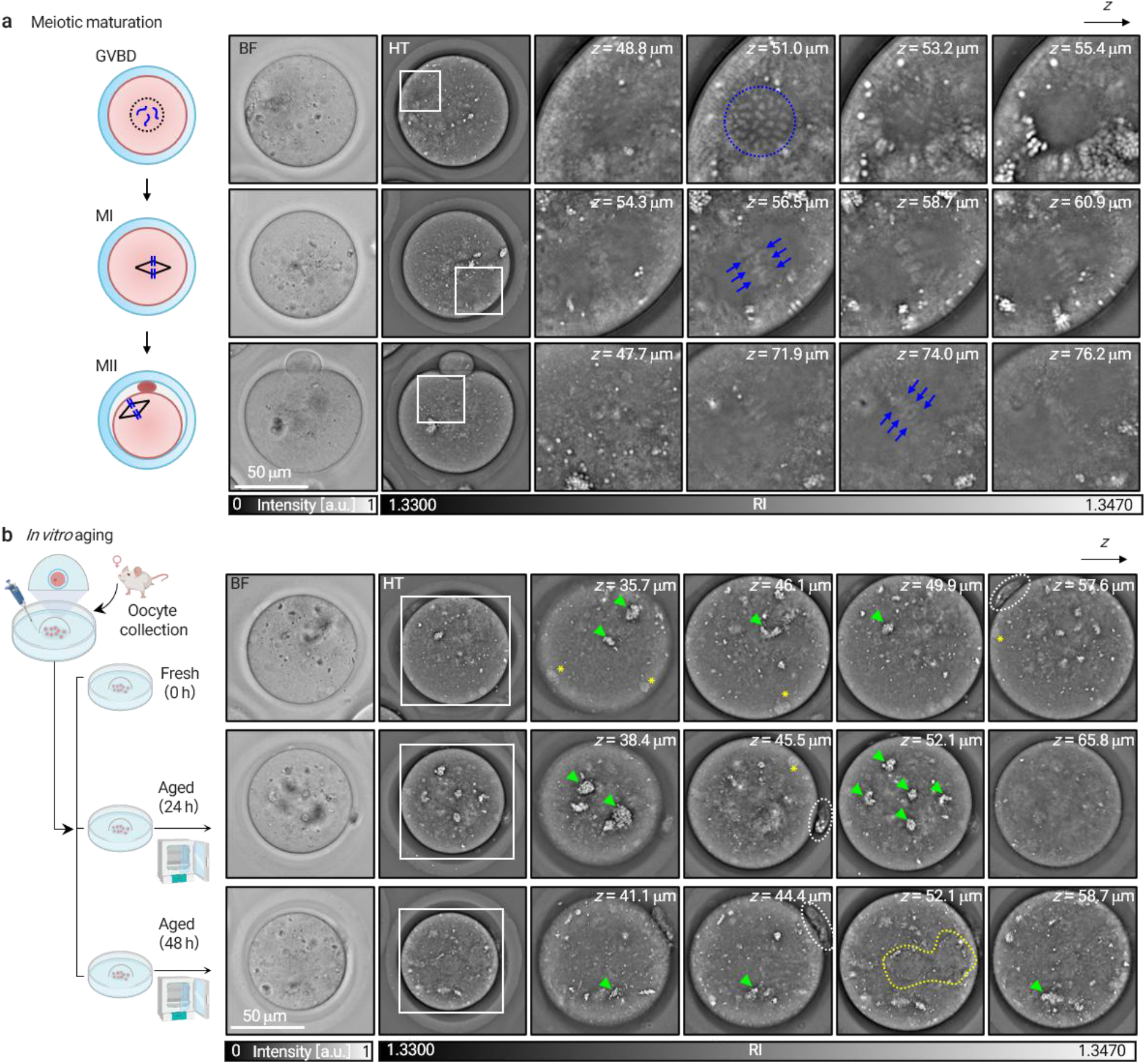
HT imaging of mouse oocytes across physiological states. **a**, Representative BF and HT images of germinal vesicle breakdown (GVBD), metaphase I (MI), and metaphase II (MII) oocytes, with axial RI slices at different depths reveal stage-associated intracellular RI organization and subcellular heterogeneity. Blue dashed circles and arrows mark RI-dense subcellular structures consistent with chromatin-associated regions. **b**, Representative BF and HT images of fresh (0 h), aged-24 h, and aged-48 h MII oocytes. Each row shows a different representative oocyte from the corresponding experimental group. Axial RI slices at different depths illustrate condition-associated differences in intracellular RI organization. Green arrowheads mark RI-dense intracellular structures; white dashed outlines mark PB regions; yellow asterisks mark peripheral RI-dense features near the oocyte cortex; yellow dashed outlines mark regions with irregular RI texture.

### Sparse-annotation AI segmentation enables compartment-resolved oocyte profiling

To convert these qualitative RI observations into compartment-resolved biophysical profiles, we next developed and validated an AI-assisted segmentation pipeline. Compartment-level quantification from 3D RI tomograms requires accurate delineation of structurally distinct regions. The pipeline combines sparse manual annotation of 5−11 representative orthogonal slices per oocyte with seeded SAM 2 video-predictor propagation across the RI stack, followed by post-processing to generate volumetric segmentations (**Fig. 4a**; see Methods). This sparse-annotation strategy reduced manual labelling burden while preserving volumetric compartment information, making large-scale phenotyping practically feasible. The predicted segmentations closely matched the underlying HT images and delineated the ooplasm, PVS, ZP, and PB in both axial and sagittal orthogonal views (**Fig. 4b**). Zoomed views across selected z-depths showed that the ZP-PVS-ooplasm interface (**Fig. 4b, i**) and PB **(Fig. 4b, ii**) were segmented consistently throughout the volume.

**Fig. 4.**
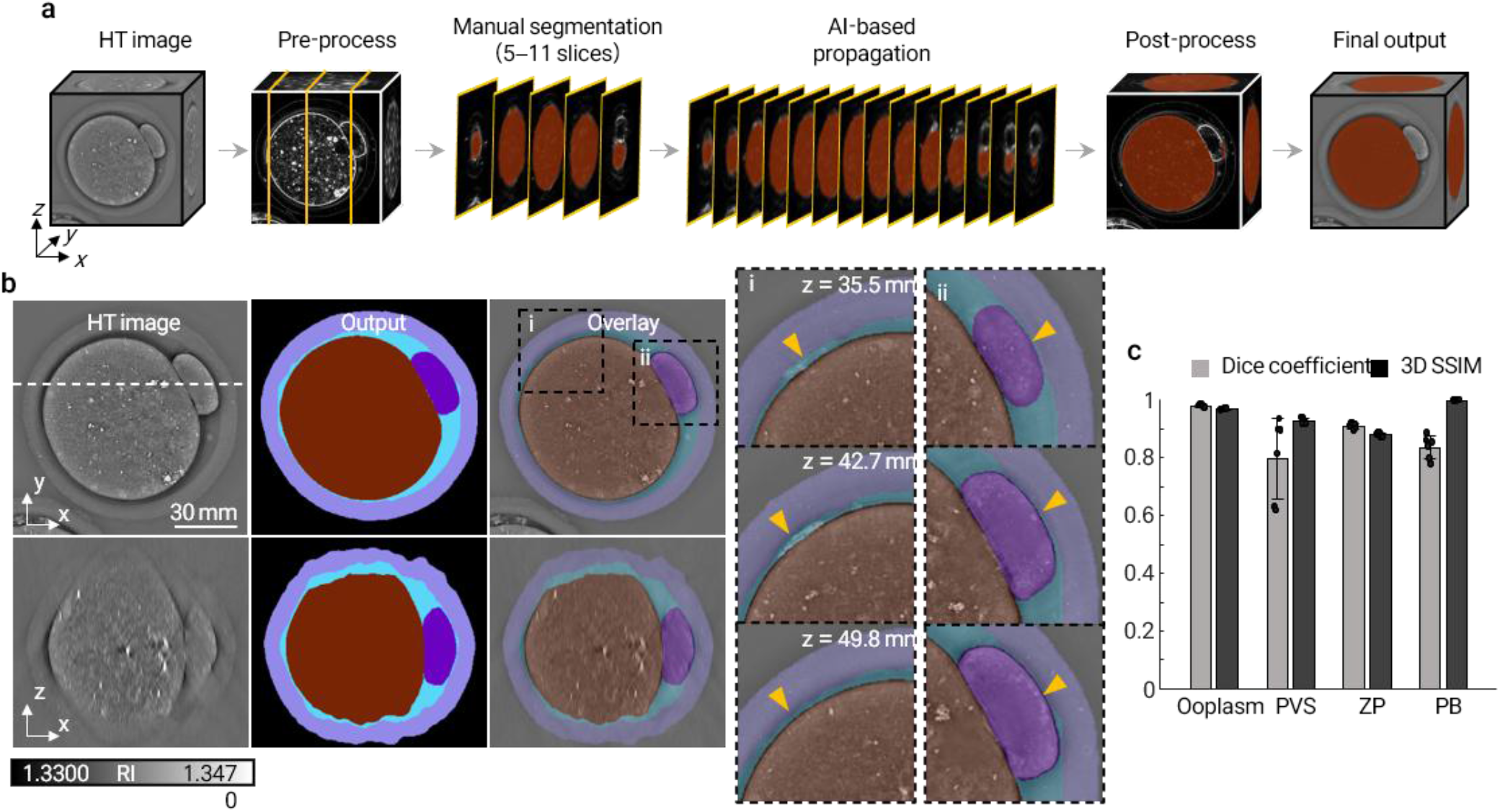
AI-assisted segmentation pipeline for 3D oocyte compartment analysis. **a**, Segmentation pipeline. Starting from the reconstructed 3D RI tomogram, a small number of representative orthogonal slices (5−11 per oocyte) are manually annotated and used as prompts for SAM 2 video-predictor propagation across the full RI stack. After post-processing, propagated per-slice masks are assembled into the final 3D compartment volume. Compartments are shown as ooplasm, brown; PVS, teal; ZP, violet; PB, dark purple. **b**, Representative final segmentation of a single oocyte. Rows show axial (XY, top) and sagittal (XZ, bottom) views; the white dashed line in the XY HT image marks the position of the XZ slice. Columns show, from left to right, the original HT image, segmentation mask, and HT-segmentation overlay. Dashed boxes in the XY overlay mark the regions shown in the zoomed views (i, ii). Insets show the ZP-PVS-ooplasm interface (i) and PB region (ii) across four z-depths (z = 35.5, 42.7, 49.8, and 58.2 μm); yellow arrowheads mark compartment boundaries and illustrate segmentation continuity across depth. **c**, Quantitative validation of segmentation performance against fully manually segmented ground-truth annotations. Dice coefficient (light grey) and three-dimensional structural similarity index measure (3D SSIM; dark grey) are shown for each compartment. Error bars indicate mean ± standard deviation (SD); points indicate individual oocytes (*n* = 6).

Quantitative validation against manual reference annotations yielded high Dice coefficients and 3D structural similarity index measure (3D SSIM) values across compartments (**Fig. 4c**), supporting the fidelity of the volumetric segmentations. Agreement was highest for the ooplasm (Dice = 0.979 ± 0.005; 3D SSIM = 0.968 ± 0.002). ZP segmentation yielded Dice = 0.910 ± 0.010 and 3D SSIM = 0.879 ± 0.007; PB segmentation yielded Dice = 0.830 ± 0.040 and 3D SSIM = 0.998 ± 0.001; and PVS segmentation yielded Dice = 0.798 ± 0.140 and 3D SSIM = 0.928 ± 0.001. These validation results indicate reliable segmentation across the major oocyte compartments and support compartment-level visualization and RI-derived feature extraction for downstream analysis.

### Compartment-resolved biophysical profiles of individual oocytes

With compartment-level segmentation in hand, we constructed a compartment-resolved, multidimensional biophysical feature space in which every oocyte is represented by 160 features: 3 RI statistics, 2 mass-related, and 16 Haralick texture features^27–29^ for each compartment, together with compartment-specific morphology features (**Fig. 5a;** see **Supplementary Note S1**). Each oocyte was therefore described not only by 2D morphology but by an integrated biophysical profile derived from its 3D RI tomogram. To introduce the feature classes that define this space, we show one representative oocyte per group: Z-score-normalized radar profiles of representative fresh, aged-24 h, and aged-48 h group oocytes illustrate the structure of this high-dimensional feature space and the position of individual oocytes within it (**Fig. 5b**).

**Fig. 5.**
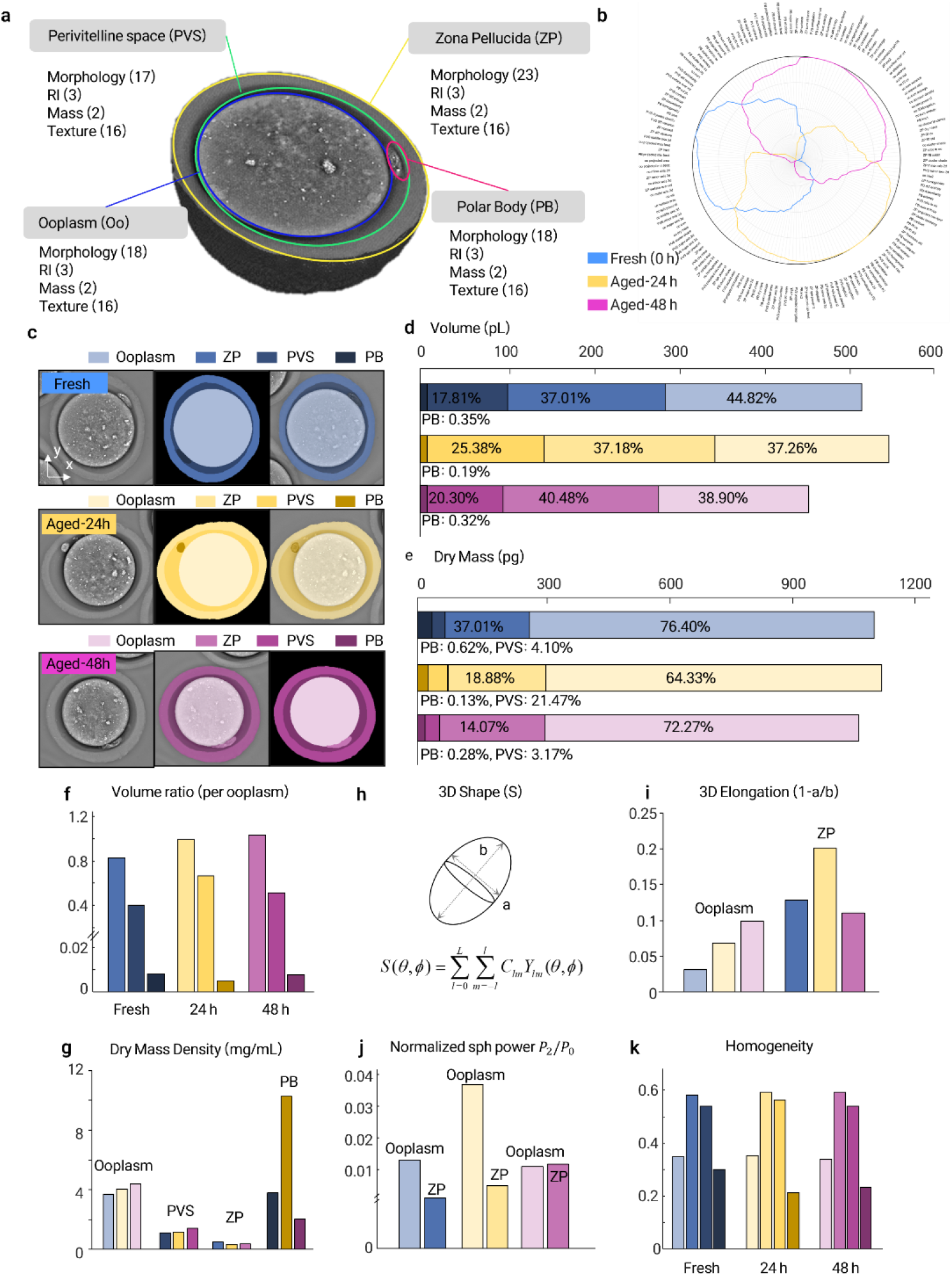
Compartment-resolved biophysical profiles of representative mouse oocytes across post-ovulatory aging conditions. **a**, Feature space extracted from each oocyte. For each of the four structural compartments — ooplasm, PVS, ZP, and PB — biophysical features were computed across four families: morphology, RI statistics, dry-mass-related quantities and Haralick texture. The full feature set comprised 160 features per oocyte; feature definitions are provided in Supplementary Note S1. **b**, Z-score-normalized radar profiles of the three representative oocytes shown in **c**. Each axis corresponds to one feature. **c**, Representative fresh, aged-24 h and aged-48 h oocytes. Columns show, from left to right, the in-focus HT image, segmentation mask, and HT–segmentation overlay, with compartments shown using the group-specific colour palette indicated above each row. **d, e**, Compartment volume and dry-mass composition of the representative oocytes. Stacked bars show total volume or dry mass, with percentages indicating compartment fractions. **f**, Ooplasm-normalized compartment volume ratios for ZP, PVS, and PB. **g**, Compartment-level dry mass density. **h**, Spherical-harmonic shape descriptor used to quantify 3D shape; the ellipsoid schematic indicates the major and minor principal axes used to compute 3D elongation. **i**, 3D elongation of the ooplasm and ZP. **j**, Normalized second-order spherical-harmonic power, *P*_2_, of the ooplasm and ZP. **k**, Haralick homogeneity of ooplasm, ZP, PVS, and PB.

Compartment-level geometric and mass-related readouts illustrate how individual feature families populate this space. Volumetric RI reconstructions of representative fresh, aged-24 h, and aged-48 h group oocytes delineated the ooplasm, ZP, PVS, and PB (**Fig. 5c**), yielding absolute compartment volume, dry mass, ooplasm-normalized volume ratios, and dry mass density (**Fig. 5d–g**). These RI-derived mass readouts provide information beyond conventional BF morphology. 3D shape was characterized by 3D elongation, 1 - a/b, and by normalized second-order spherical-harmonic power, *P*_2_ (**Figs. 5h–j**; see Methods). Finally, intracellular Haralick texture, illustrated here by homogeneity, captured spatial organization of RI signals within the ooplasm (**Fig. 5k;** see Methods). Together, these representative profiles show how HT integrates volumetric morphology, mass, shape, and intracellular texture into a unified biophysical description of individual oocytes.

### Coordinated multidimensional remodeling across post-ovulatory aging conditions

Having introduced the compartment-resolved biophysical feature space through representative oocytes (**Fig. 5**), we next asked whether ooplasm-derived features distinguish the three post-ovulatory aging states at the population level across the full MII cohort (fresh, *n* = 96; aged-24 h, *n* = 77; aged-48 h, *n* = 49; **Fig. 6a**). We focused on the ooplasm because it is the primary compartment undergoing cytoplasmic reorganization during *in vitro* post-ovulatory aging^25,30^ and showed the most robust segmentation performance among compartments.

**Fig. 6.**
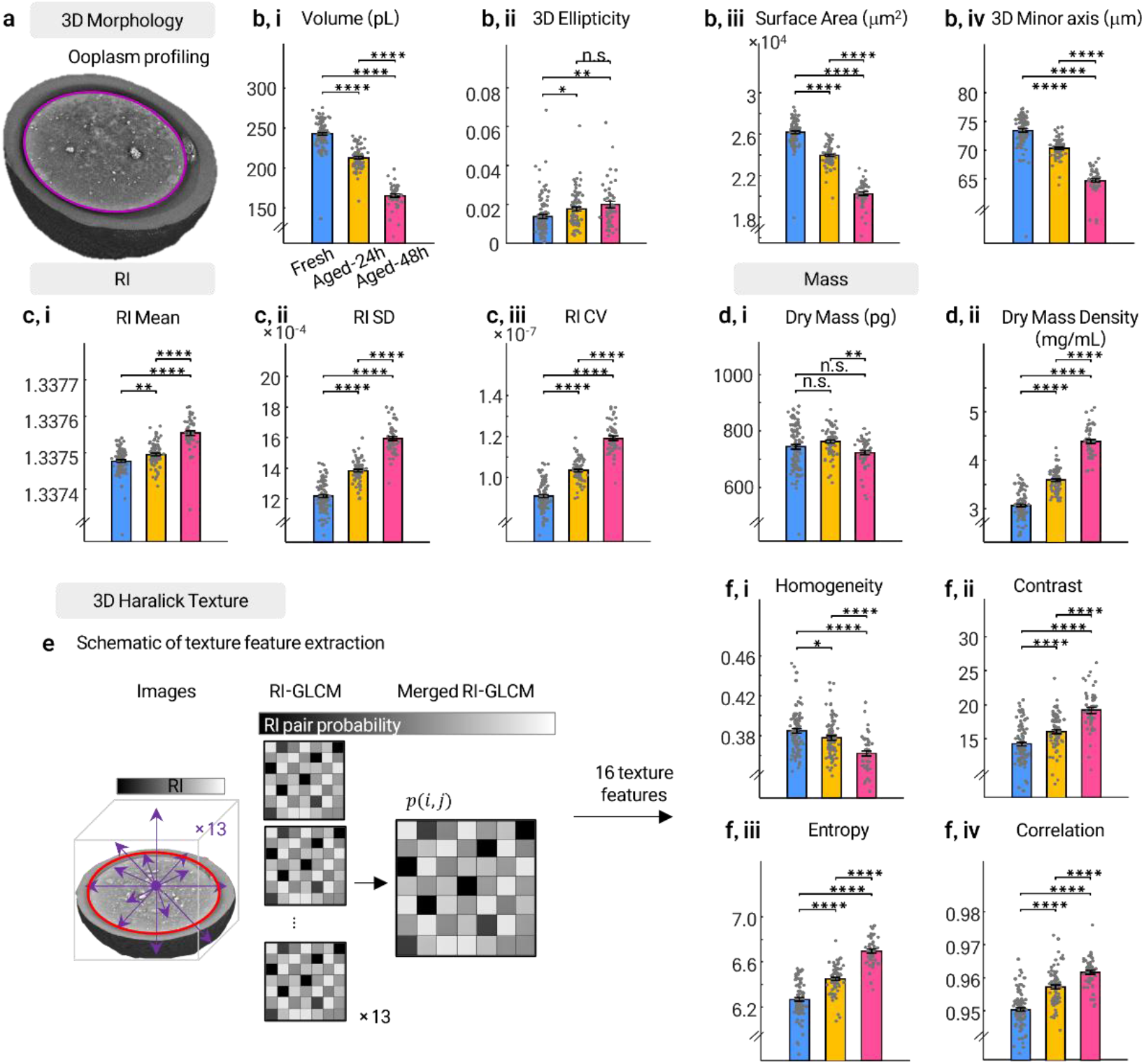
Population-level ooplasm biophysical features distinguish fresh and aged mouse oocytes. **a**, Representative 3D RI rendering of an MII mouse oocyte; the yellow contour marks the ooplasm boundary used for ooplasm-level feature extraction in **b**–**f. b**, 3D ooplasm morphology features: volume (i), 3D ellipticity (ii), surface area (iii), 3D minor-axis length (iv). **c**, Ooplasm RI statistics: mean (i), standard deviation (SD) (ii), and coefficient of variation (CV) of RI values. **d**, Ooplasm dry-mass-related features: total dry mass (i) and dry-mass density (ii). **e**, Schematic for 3D Haralick texture extraction. Thirteen symmetry-equivalent 3D directions (violet arrows) generate per-direction grey-level co-occurrence matrices (GLCMs, *p*(*i, j*)) over the ooplasm volume; these are merged into an orientation-averaged GLCM from which 16 Haralick texture features are computed. **f**, Representative 3D Haralick texture features of the ooplasm: homogeneity (i), contrast (ii), entropy (iii), and correlation (iv). Feature distributions are shown for fresh (0 h; *n* = 96), aged-24 h (*n* = 77), and aged-48 h (*n* = 49) MII oocytes. Data were collected across three independent experiments. Bars indicate mean ± SD; dots indicate individual oocytes. Colour code: fresh, blue; aged-24 h, yellow; aged-48 h, magenta. For each of the 39 ooplasm features, between-group comparisons were performed using one-way ANOVA followed by Tukey’s post hoc tests (18 morphology + 3 RI + 2 mass + 16 Haralick texture). Significance levels are indicated as *p* < 0.05(*), *p* < 0.01(**), *p* < 0.001(***), *p* < 0.0001(****); n.s., not significant.

3D ooplasm morphology differed across the three conditions at the population level (**Fig. 6b;** expanded 14-panel morphology breakdown in **Supplementary Fig. S2**). Ooplasm volume decreased monotonically from fresh to aged-24 h and aged-48 h oocytes (fresh: 242.80 ± 18.05 pL; aged-24 h: 212.85 ± 13.48 pL; aged-48 h: 165.59 ± 14.67 pL; Fig. 6b, i). 3D ellipticity increased modestly with aging but did not differ significantly between aged-24 h and aged-48 h groups (fresh: 0.014 ± 0.010; aged-24 h: 0.018 ± 0.010; aged-48 h: 0.020 ± 0.011; **Fig. 6b, ii**). Surface area showed a parallel decrease across the same groups (fresh: (26.20 ± 1.33) × 10^3^; aged-24 h: (24.00 ± 1.06) × 10^3^; aged-48 h: (20.30 ± 1.20) × 10^3^ μm^2^; Fig. 6b, iii), as did 3D minor-axis length (fresh: 73.42 ± 2.87 μm; aged-24 h: 70.31 ± 1.63 μm; aged-48 h: 64.73 ± 2.29 μm; Fig. 6b, iv). Together, these morphological metrics were consistent with coordinated ooplasm shrinkage across aging conditions.

In parallel, ooplasm RI statistics, mean RI, RI SD, and RI coefficient of variation (CV), shifted progressively with aging (**Fig. 6c**), indicating age-associated redistribution of intracellular RI within the ooplasm, a trend that was further reflected in the texture-level heterogeneity described below. Mean RI increased across the three groups, from 1.3374 ± 0.36 × 10^−4^ in fresh oocytes to1.3375 ± 0.33 × 10^−4^ at aged-24 h and 1.3376 ± 0.46 × 10^−4^ at aged-48 h. RI SD (fresh: 12.18 ± 0.99 × 10^−4^; aged-24 h: 13.84 ± 1.71 × 10^−4^; aged-48 h: 15.94 ± 1.04 × 10^−4^), and RI CV (fresh: 9.11 ± 0.74 × 10^−7^; aged-24 h: 10.34 ± 0.52 × 10^−7^; aged-48 h: 11.91 ± 0.78 × 10^−7^) increased monotonically across every pairwise comparison, reflecting a progressive increase in intracellular RI magnitude together with a monotonic rise in intracellular RI heterogeneity. Notably, unlike the intensity-based analysis (see **Supplementary Fig. S3**), where the mean plateaued between aged-24 h and aged-48 h, the RI mean continued to rise between these time points; this mean shift is small, however, relative to the monotonic increases in RI heterogeneity (SD and CV) that dominate the aging signal.

Dry mass-related descriptors showed that ooplasm shrinkage was accompanied by increased dry-mass concentration rather than a proportional decrease in ooplasm dry mass (**Fig. 6d**). Total ooplasm dry mass remained broadly conserved, with group means of 743.92 ± 81.57, 762.39 ± 47.34, and 723.31 ± 52.97 pg in fresh, aged-24 h, and aged-48 h oocytes, respectively. In contrast, ooplasm dry-mass density rose monotonically across the same groups, from 3.07 ± 0.27 to 3.59 ± 0.24 and 4.38 ±0.31 mg/mL, respectively (**Fig. 6d, ii**), whereas total ooplasm dry mass was statistically indistinguishable between fresh and aged-24 h oocytes and only modestly lower at aged-48 h (**Fig. 6d, i**). The progressive volume reduction (**Fig. 6b, i**), combined with conserved-to-modestly-declining total dry mass (**Fig. 6d, i**) and monotonically rising dry-mass density (**Fig. 6d, ii**), is operationally consistent with cytoplasmic condensation rather than net mass loss, a readout accessible because label-free HT returns absolute RI from which dry mass can be estimated under a standard RI-increment assumption (Methods).

Intracellular Haralick texture descriptors (homogeneity, contrast, entropy, and correlation) shifted progressively across aging conditions (**Fig. 6e, f;** full 16-panel ooplasm texture breakdown in Supplementary **Fig. S4**). Homogeneity decreased monotonically while contrast, entropy, and correlation all increased monotonically, indicating a progressive loss of local RI uniformity and a rise in intracellular RI heterogeneity that is invisible to morphological analysis alone. These population-level results indicate coordinated aging-associated differences in ooplasm morphology, mass distribution, and intracellular spatial organization, motivating a direct benchmark of the HT feature panel against conventional BF analysis.

### HT-derived biophysical profiles strengthen aging-state separation relative to brightfield-derived profiles

To benchmark the value added by holotomography information, we compared feature sets derived from single in-focus BF images (2D BF), multi-slice BF stacks (2.5D BF), and volumetric HT data (HT; **Fig. 7a**). The BF feature sets were constructed from 2D descriptors reported in the literature^5^ (see Supplementary Note S2), including conventional morphometry, intensity statistics, and grey-level co-occurrence matrix (GLCM) texture features computed directly from BF intensity images^21^. The HT feature set built on this foundation by incorporating related descriptor classes together with information available from volumetric RI data, including 3D morphology, RI statistics, and dry-mass-related features (see Supplementary Note S3). Uniform manifold approximation and projection (UMAP) embeddings showed progressively clearer separation of fresh, aged-24 h, and aged-48 h oocytes from 2D BF to 2.5D BF and HT, using *n*_neighbors = 15 and min_dist = 0.10.

**Fig. 7.**
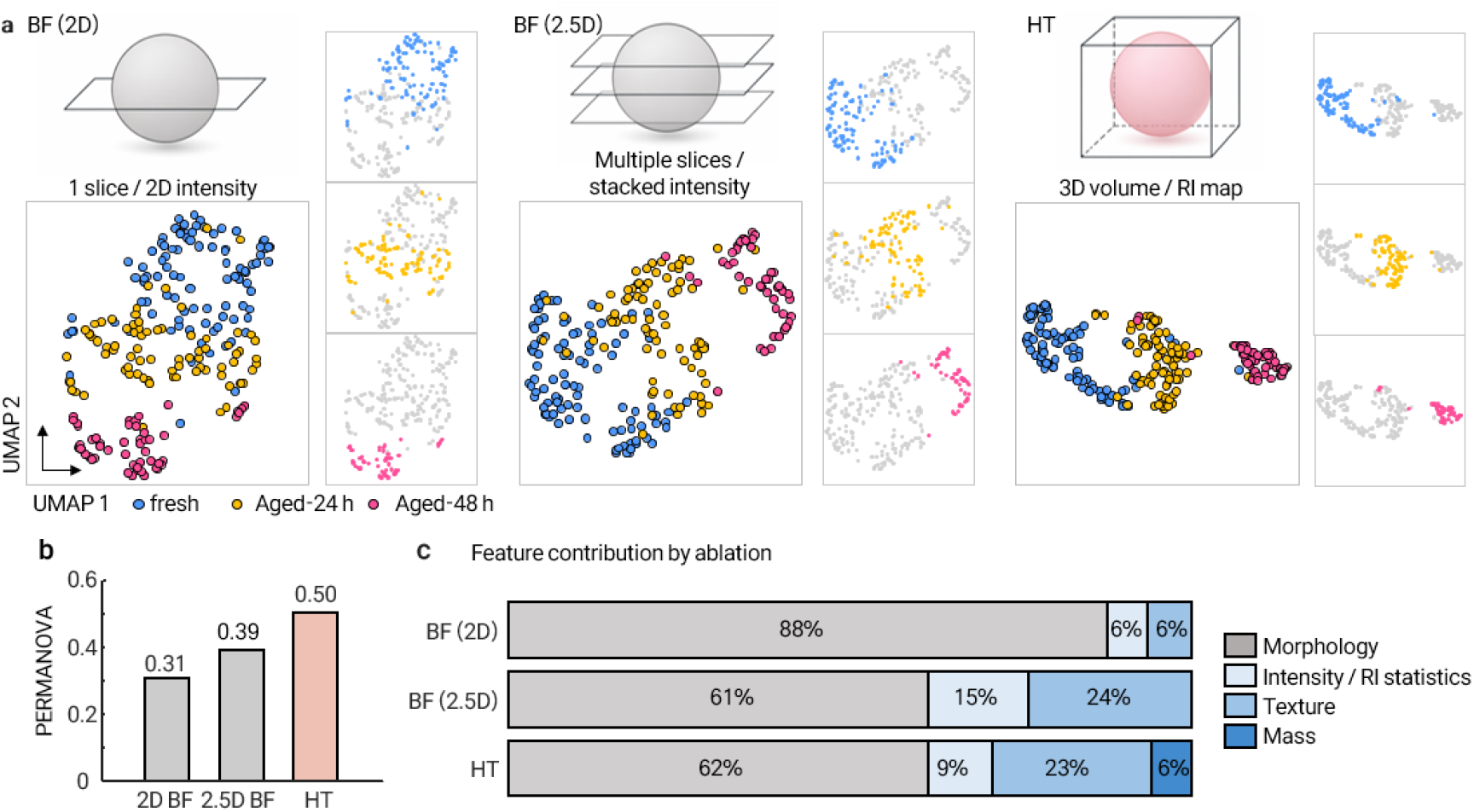
Comparison of aging-state separation across brightfield and HT-derived feature sets. **a**, Feature extraction and UMAP embedding from brightfield (2D BF), multi-slice brightfield (2.5D BF), and HT data. 2D BF features were extracted from a single in-focus intensity image; 2.5D BF features were extracted from 11 intensity images sampled at 15 µ m axial intervals across the oocyte volume; HT features were extracted from the full 3D RI volume. Points represent individual MII oocytes from the fresh (0 h, *n* = 96; blue), aged-24 h (*n* = 77; yellow) and aged-48 h groups (*n* = 49; pink). Data were collected across three independent experiments. **b**, Group-separation performance quantified by PERMANOVA *R*^*2*^ for 2D BF, 2.5D BF and HT feature sets. **c**, Feature-family contribution to group separation by ablation. Contributions were calculated from leave-one-feature-out *R*^*2*^ values and are shown as relative contributions.

To quantify the separation observed in the UMAP embeddings, we evaluated each feature set using permutational multivariate analysis of variance (PERMANOVA) and silhouette scores. PERMANOVA *R*^*2*^ increased from 0.31 for 2D BF to 0.39 for 2.5D BF and 0.50 for HT (**Fig. 7b**). Silhouette scores showed the same trend, increasing from 0.18 to 0.24 and 0.35, respectively (see Supplementary **Fig. S7**). Thus, features derived from volumetric RI data provided stronger population-level separation of fresh, aged-24 h, and aged-48 h oocytes than features derived from BF images.

We next asked which feature families contribute to this separation. We ablated feature families in turn and measured the resulting loss in group separation using the PERMANOVA score (**Fig. 7c**). Features whose removal improved group separation were excluded from the contribution calculation, because their removal improving separation indicates they added noise to the embedding rather than discriminative signal; the reported percentages therefore reflect the relative contributions of features that supported separation. In 2D BF, separation was dominated by morphology-related features (88%), with intensity and texture descriptors accounting for 12%. In contrast, morphology-related features accounted for 62% of HT separation, whereas RI statistics, dry-mass-related features, and texture together accounted for 38%. Considering that the absolute number of morphology features in HT (18 features) is twice that of BF (9 features), these results indicate that the improved separation in the HT feature space was supported substantially by RI-derived, non-morphological descriptors rather than by morphology alone.

We further performed a leave-one-feature-out ablation analysis on the silhouette score, with the expanded feature-level ΔSil ranking shown in **Supplementary Fig. S7**. Features whose removal improved silhouette-based group separation were excluded from this ranking, so the analysis highlights individual features that supported separation.

## Discussion

The central finding of this work is that label-free 3D RI imaging supports mouse oocyte phenotyping not only through volumetric information, but also through quantitative biophysical descriptors derived from RI. Ablation analysis of 2D BF, 2.5D BF, and HT feature sets showed that non-morphological descriptors (dry mass, intracellular texture, and RI statistics) collectively accounted for 38% of HT-based separation between aging states, whereas 2D BF separation was dominated by morphology-related features (88%). This partition is a descriptive decomposition of feature-set contributions to population-level separation rather than a mechanistic isolation of independent biophysical axes, since RI statistics, dry-mass density, and intracellular texture are not fully orthogonal to ooplasm volume. The advantage of HT therefore does not appear to arise simply from increased dimensionality, but from access to RI-derived feature classes that are fundamentally inaccessible to BF microscopy: dry-mass density, absolute RI statistics, and volumetric RI-based texture require direct measurement of the RI field and cannot be recovered from intensity images alone. Among reported label-free implementations applied to mouse oocytes, HT provides a complementary axis: it is the only one summarized here that yields volumetric RI information from which dry mass, RI statistics, and intracellular texture can be derived. By contrast, back-scattering modalities, including optical coherence microscopy, optical coherence tomography, and reflection-matrix microscopy, provide structural contrast. Nonlinear and vibrational modalities, including higher-harmonic generation, CARS and Raman microscopy, probe nonlinear or chemical contrast; and Brillouin microscopy probes mechanical properties (Supplementary Table 1)^6,9,11,31^. Volumetric RI therefore provides a complementary analytical axis for label-free oocyte phenotyping. Parallel work using quantitative phase imaging has demonstrated dry-mass-based characterization of embryo components and reproductive outcome prediction^32^, and digital holographic microscopy has been used to follow embryo RI dynamics^33^; broader reviews of label-free holographic imaging in gametes and embryos have surveyed the rapidly evolving landscape^34–36^.

The framework also reveals two biophysical observations about mouse oocytes imaged in our in vitro model of post-ovulatory aging^25,26^. First, in vitro post-ovulatory exposure was associated with coordinated differences across multiple ooplasm feature classes — morphology, mass, RI, and texture (**Fig. 6**), rather than a shift in any single parameter. Second, aged oocytes displayed an RI-derived signature in which dry-mass density increased while ooplasm volume decreased and total integrated dry mass was conserved or only modestly reduced (**Fig. 6d**), a combination consistent with volume reduction accompanied by intracellular compaction rather than bulk mass loss. We do not attach a specific mechanism to this signature. The observed pattern is compatible with multiple processes, including cytoplasmic reorganization associated with actin-coated vesicle motility^37^, mitochondrial clustering and endoplasmic reticulum redistribution under post-ovulatory oxidative stress^25,26,38^, cortical granule loss^39^, and possible cytoplasmic dehydration during culture. Of these, cytoplasmic dehydration and organelle clustering are most directly consistent with the observed combination of increased density and conserved integrated mass, although our data cannot adjudicate among the candidate processes. Correlative imaging with organelle-specific markers, such as MitoTracker, ER-Tracker, or cortical granule probes, will be required to assess their relative contributions^30,40–42^.

Interpretation of individual RI-dense features also requires caution. RI contrast is sensitive to local composition and density but does not assign molecular identity. Qualitative spatial co-localization on a small number of selected oocytes between HT contrast and fluorescence signals from Hoechst and LipiDye II (Fig. 2) supports the biological relevance of the RI maps without establishing one-to-one correspondence between RI features and molecular components. Chemically specific modalities such as CARS and Raman are better suited for assigning molecular composition to individual intracellular structures^31,43^. Changes in texture, RI variance, and mass-related properties are consistent with intracellular remodeling, but their mechanistic basis remains to be defined. Correlative studies combining HT with fluorescence markers, ultrastructural imaging, or perturbation experiments will be valuable for linking specific RI-derived phenotypes to underlying cellular processes.

The principal translational value of this work lies not in characterizing in vitro post-ovulatory aging itself, but in establishing that a clinically relevant axis of oocyte quality exists below the detection threshold of conventional microscopy and can be quantified by label-free 3D RI imaging. In current clinical IVF practice, MII oocyte selection relies almost entirely on BF morphological assessment, yet morphologically normal MII oocytes routinely differ in fertilization rate, blastocyst formation, and euploidy — pointing to a dimension of oocyte quality not captured by conventional brightfield grading. The in vitro aging paradigm employed here serves as an experimentally amplified system in which such deterioration is deliberately induced and made detectable: the HT-derived features that diverged between fresh and aged oocytes (dry-mass density, RI heterogeneity, and 3D texture entropy) fall precisely within the feature class that BF imaging cannot recover, establishing proof-of-concept that biophysical quality information beyond the BF detection limit is both real and quantifiable^44–48^.

A directly testable hypothesis follows: the same RI-derived axes may resolve quality differences among freshly retrieved, morphologically normal MII oocytes. Dry-mass density and RI heterogeneity are the most plausible translation candidates, as they probe cytoplasmic compaction and organizational disorder mechanistically tied to mitochondrial distribution and cytoplasmic maturation, whereas 3D texture entropy provides a complementary, geometry-free descriptor of intracellular order. Two caveats temper this extrapolation. First, the biophysical signatures of aging-induced deterioration and of incomplete cytoplasmic maturation may not be identical and could, for some features, point in opposite directions, dry-mass density, for example, may rise with dehydration but fall with incomplete maturation. Second, the dynamic range of HT signals between fresh and aged oocytes is amplified by design, and inter-oocyte variation among fresh MII oocytes will be subtler and require correspondingly higher measurement precision. Both points define empirical questions rather than barriers in principle. The natural next step is therefore a prospective study in which freshly retrieved MII oocytes are imaged non-invasively by HT prior to insemination and followed through fertilization, blastocyst formation, and, where feasible, euploidy and live-birth outcomes^49,50^.

Four limitations bound the present work and define near-term extensions.

i. Model and species scope. The aging axis studied here is an in vitro model — MII oocytes cultured for 24 h or 48 h in M16 medium under mineral oil at 37°C in 5% CO_2_, and therefore reflects a specific combination of culture-induced oxidative, metabolic, and osmotic stresses rather than physiological reproductive aging in the oviduct^25,26^. The RI-derived signatures reported here should be interpreted as biophysical consequences of in vitro post-ovulatory exposure. Their correspondence with in vivo aging, such as oocytes from reproductively aged females or oocytes retrieved after delayed mating, and with antioxidant-rescue controls remains to be established. Experiments were also performed exclusively in mouse oocytes, and extension to human oocytes will be essential for translational relevance. Feature-specific generalizability may vary: compartment-level RI statistics, dry-mass density, and Haralick texture are likely to be more transferable, whereas absolute geometric descriptors, including ZP thickness, PVS volume, and spherical-harmonic coefficient magnitudes, will require species-specific recalibration in matched human-oocyte cohorts.
ii. Developmental competence not assayed. We did not assay developmental competence (fertilization, cleavage, or blastocyst formation) for the imaged oocytes; the present features therefore characterize an in vitro post-ovulatory aging state rather than predict developmental potential per se. Evaluating the biomarker potential of these signatures for developmental competence will require prospective studies in which MII oocytes are imaged non-invasively before insemination and followed through fertilization and embryo development. Recent BF deep-learning approaches have shown that intensity images alone can predict chromatin state and developmental potential in live mouse oocytes^51^. Such approaches, however, extract latent features whose biophysical meaning is not directly interpretable; volumetric RI is complementary in providing explicit, quantitative, and platform-independent access to dry mass and 3D texture, and combining the two modalities is a natural next step^52,53^.
iii. Compartment-level quantification. Compartment-level analysis is bounded by segmentation accuracy and by the ooplasm-focused scope of the present work. Although the pipeline enabled whole-oocyte compartment visualization, population-level statistical analyses focused on the ooplasm, which showed the most robust segmentation performance. Quantitative interpretation of ZP-, PVS-, and PB-derived descriptors therefore remains preliminary, particularly for the thin and variable PVS, and should be extended to population-level analysis in future work.
iv. Reconstruction and instrument dependence. Absolute RI-derived values, including dry-mass density and Haralick descriptors, may depend on the reconstruction algorithm and the optical system. Patterned-illumination HT has anisotropic spatial support, with coarser axial than lateral resolution owing to the missing cone of the illumination numerical aperture. Shape descriptors that combine lateral and axial extent, particularly *P*_2_ and major-axis length, may therefore inherit an instrument-specific baseline. The same anisotropic support and iterative regularization also influence axially resolved RI gradients, and therefore the absolute magnitudes of RI statistics (SD, CV) and of volumetric Haralick texture, which are computed over the reconstructed volume; because axial optical resolution is coarser than lateral, orientation-averaged 3D texture is dominated by lateral structure and should be read as a platform-native readout. Because the aging comparisons here were performed between groups imaged on the same platform, the relative trends should be robust to this baseline. However, inter-laboratory and inter-instrument validation will be required for cross-platform deployment and for absolute value reporting.

Biomedical imaging has long relied on appearance-based assessment of cells, tissues, and organs, despite growing evidence that biophysical state shapes function. The work reported here illustrates how label-free volumetric imaging, compartment-resolved segmentation, and ablation-based feature decomposition can translate qualitative morphology into quantitative biophysical profiling of intact mammalian oocytes. The same approach can in principle be extended to embryos, organoids, and thick tissue sections, specimens where destructive labelling is not an option, subject to the inter-instrument validation noted above. By shifting the analysis from appearance alone to quantitative biophysical composition and spatial organization, label-free 3D phenotyping provides a quantitative framework for non-invasive characterization of aging-associated biophysical state in mammalian oocytes, with potential extension to embryos, organoids, and other intact specimens where destructive labelling is not an option.

## Methods

### Experimental design

The study establishes a label-free 3D biophysical phenotyping framework for mouse oocytes using HT. We collected intact oocytes, grouped them by in vitro post-ovulatory aging condition, and for each oocyte acquired a label-free holotomography volume, reconstructed its 3D RI distribution, segmented its major compartments (ooplasm, ZP, PVS, and PB) and extracted morphology-based and RI-derived biophysical features for comparative analysis. For compartment-wise single-oocyte comparison, we extracted a total of 160 HT-derived features across the segmented oocyte compartments.

For population-level statistical analysis, we focused on the ooplasm and compared aging-associated feature distributions across the three MII oocyte groups: fresh, aged-24 h, aged-48 h. Ooplasm-level features were organized into four categories — morphology, RI/intensity statistics, Haralick texture, and dry mass — to quantify category-specific remodeling across aging conditions. For this analysis, we used 39 HT-derived ooplasm features (18 morphology + 3 RI + 2 mass + 16 Haralick texture). We additionally computed matched 2D and multi-slice (2.5D) BF feature sets from the same samples, each comprising 28 features (9 morphology + 3 intensity + 16 Haralick texture), to assess the information added by volumetric RI imaging.

Feature selection differed between the two analyses because they served distinct purposes. The compartment-wise single-oocyte analysis was designed to maximize the descriptive feature space across oocyte compartments, whereas the population-level statistical analysis was designed for modality-balanced comparison across HT, 2D BF, and 2.5D BF.

### Animals and ethics statement

All animal procedures were approved by the Institutional Animal Care and Use Committee (IACUC) of the Korea Advanced Institute of Science and Technology (KAIST; protocol KA2024-021-v2) and followed relevant institutional and national guidelines. Female C57BL/6J mice aged 4 weeks were maintained at the KAIST Laboratory Animal Resource Center under standard husbandry conditions, with ad libitum access to standard rodent chow and reverse-osmosis water. Five female mice were used per experimental replicate, and the complete experimental cycle was repeated three times.

### Mouse oocyte collection and classification

Cumulus–oocyte complexes (COCs) were collected from the oviductal ampullae following superovulation. Each female received an intraperitoneal injection of 0.1 mL CARD HyperOva (Cosmo Bio, Japan), followed 48 h later by an intraperitoneal injection of 20 IU human chorionic gonadotropin (hCG; Sigma-Aldrich, USA). At 15−18 h post-hCG, animals were euthanized by cervical dislocation and the oviducts were excised. COCs were washed in 80 μL drops of human tubal fluid (HTF; Cosmo Bio, Japan) under light mineral oil (GeorgiaChem, USA) pre-equilibrated overnight at 37°C in 5% CO_2_. Oocytes were classified as GVBD, MI, or MII based on external morphology by an experienced embryologist; oocytes with overt morphological abnormality at the time of imaging (e.g., parthenogenetic activation, missing ZP, pronounced fragmentation) were excluded from downstream analysis. Across the three independent experimental cycles, exclusions were rare (<5% of recovered MII oocytes per group), and imaging order across the fresh, aged-24 h, and aged-48 h groups was randomized within each replicate. Because collection at 15−18 h post-hCG yields a cohort dominated by MII oocytes with a minority of GVBD and MI oocytes, quantitative population-level analyses of post-ovulatory aging were restricted to the MII cohort; GVBD and MI oocytes are presented in **Fig. 3a** as imaging-level demonstrations of state-specific intracellular organization. For aging analysis, freshly collected MII oocytes formed the fresh group (0 h; see *In vitro post-ovulatory aging* for the operational definition of 0 h), and additional MII oocytes were maintained in vitro and imaged after 24 h or 48 h of culture (denoted fresh, aged-24 h, and aged-48 h). Oocytes were handled to preserve viability and structural integrity throughout sample preparation and imaging.

### In vitro post-ovulatory aging

Immediately after collection, cumulus-oocyte complexes (COCs) were denuded by incubation in HTF containing 1% hyaluronidase (Sigma-Aldrich, USA) at 37°C for 1 min, followed by immediate washing in HTF. The denuded MII oocytes were then subjected to in vitro post-ovulatory aging^25,26^. Freshly collected MII oocytes were cultured in 20 μL drops of M16 medium (Merck) covered with 2 mL of mineral oil in confocal dishes, with 7−10 oocytes per drop. The culture dishes were pre-incubated for 3 h at 37°C in a humidified 5% CO_2_ atmosphere before use. Oocytes assigned to the aged groups were cultured for 24 h or 48 h in a humidified CO_2_ incubator before transfer to a Tokai Hit on-stage incubator for imaging; freshly collected oocytes imaged without prolonged culture served as the (0 h) fresh group. For each group, the 0 h reference corresponds to the moment immediately prior to image acquisition rather than to the moment of oocyte retrieval; the interval between retrieval and the onset of 0 h imaging was approximately 40–50 min, spanning transfer to the imaging dish and system set-up. Imaging of the 0 h, 24 h, and 48 h groups was performed daily between 11:00 and 13:00 over three consecutive days, and the complete experimental cycle was repeated three times. On the imaging stage, oocytes were maintained in M16 medium inside a Tokai Hit on-stage incubator at 37°C in 5% CO_2_ under humidified conditions; imaging was initiated immediately upon stage loading, with a brief 1–2 min equilibration applied when ambient vibration was detected.

### Holotomography, brightfield, fluorescence imaging, and refractive index reconstruction

Live oocytes were imaged on an HT system (HT-X1 MAX, water-immersion configuration; Tomocube, Republic of Korea) under patterned illumination. The system employs a low-coherence light source (LED, illumination wavelength of 660 nm ± 10 nm), and projects four optimized wavefront illumination patterns onto a sample at successive axial positions. The diffracted patterns from a sample are collected, from which the three-dimensional RI distribution of the sample is reconstructed via 3D deconvolution algorithms with iterative regularization^54–58^. The reconstructed RI is referenced to the surrounding M16 culture medium (*n*_*m*_ = 1.3370).

The same system was used to acquire HT, BF, and fluorescence images from the same live oocytes. Multiple raw intensity frames were acquired for volumetric RI reconstruction, and the corresponding raw intensity measurements were used as BF images for comparison with conventional optical imaging. Imaging was performed using a 24.33× water-immersion objective lens with a numerical aperture of 1.1. The lateral field of view was 320 × 320 µm. For each oocyte, a 3D RI tomogram spanning the full oocyte and surrounding ZP was reconstructed with a voxel size of 0.113 × 0.113 × 0.549 µm (*x* × *y* × *z*). Day-to-day RI stability was monitored across imaging sessions on medium-only fields and remained within the platform-level repeatability of the instrument. Reconstructed RI volumes were represented in Cartesian coordinates and visualized as orthogonal planes and selected axial slices. For downstream analysis, each volume was exported as a stack of RI images and processed with custom scripts written in Python and MATLAB. Oocytes were loaded individually into confocal dishes containing M16 medium and imaged under live, label-free conditions, with care taken to minimize motion and optical perturbation during acquisition.

For selected oocytes, fluorescence images were additionally acquired to assess the biological relevance of RI contrast. Live oocytes were incubated for 1 h at 37°C with Hoechst 33342 Ready Flow Reagent (Invitrogen, Thermo Fisher Scientific, catalog no. R37165) at 2 drops/mL to label nuclear material, or with LipiDye II lipid droplet live-imaging dye (Funakoshi, catalog no. FDV-0027) at 1 μM, prepared by 1:1000 dilution from a 1 mM stock solution, to label lipid-rich structures.

Fluorescence and BF images were registered with the corresponding HT images to generate overlay views for qualitative comparison; fluorescence imaging served only as a reference for biological correspondence and was not used for primary quantitative phenotyping.

### Oocyte compartment definition, manual annotation, and AI-assisted segmentation

We defined four structural compartments for downstream quantification from the reconstructed RI tomograms: ooplasm, ZP, PVS, and PB. The ooplasm was defined as the main cell body enclosed by the oolemma; the ZP as the extracellular glycoprotein layer surrounding the oocyte; the PVS as the region between the oolemma and ZP; and the PB as the small cellular structure located within the PVS.

These compartments were segmented by combining sparse manual annotation with AI-based propagation using a pre-trained Segment Anything Model 2 (SAM 2) foundation model operated in its video predictor mode. We used the hiera-small SAM 2.1 checkpoint and its corresponding configuration file. For each oocyte, 5−11 representative axial slices, including the center slice, were manually annotated to generate mask prompts for SAM 2-based propagation across the full preprocessed image stack. Predicted volumes were refined by distance-transform-based watershed refinement followed by dominant-region selection, compartment-specific morphological opening and closing, and Gaussian smoothing with σ = 2 voxels, producing the final compartment masks. This strategy reduced the manual labelling burden relative to full-stack annotation while preserving volumetric compartment information. Manual annotations were supplied as full per-slice mask prompts on the representative axial slices, propagated bidirectionally (forward and backward from each annotated slice) by the SAM 2 video predictor. The number of annotated slices per oocyte (5−11) varied with compartment complexity rather than aging condition, and annotators were not blinded to aging condition during the labelling step^59^.

### Segmentation validation

To generate ground-truth references for validation, a held-out subset of oocytes was fully manually segmented slice-by-slice with dense per-compartment annotation (ooplasm, 3−6 slices per oocyte; PVS, 21−34 slices; ZP, 11−26 slices; PB, 5−9 slices), where the per-compartment slice counts reflect the native axial extent of each compartment rather than sub-sampling. We evaluated segmentation performance by comparison against these ground-truth annotations, using the Dice coefficient and the 3D structural similarity index measure (3D SSIM) for each compartment (ooplasm, PVS, ZP, PB). For validation, the original RI volumes had a size of 1201 ⨯ 1201 ⨯ 896 voxels. Volumes were spatially downsampled by a factor of 0.3 in the lateral dimensions and permuted into the SAM 2 input orientation, yielding 360 2D frames per oocyte for validation. Segmentation validation was performed on six oocytes by comparing the AI-assisted segmentation results with the corresponding manual reference annotations. Population-level statistical analyses subsequently focused on the ooplasm compartment, for which segmentation performance was strongest (Dice 0.979 ± 0.005); ZP, PVS, and PB compartment-level descriptors should therefore be considered preliminary and were not used as the basis for the main population-level conclusions.

### Extraction of compartment-level morphological features

From the segmented 3D masks, we computed compartment-level morphological features. Absolute volume was calculated as the product of segmented-voxel count and voxel volume; to normalize for oocyte size, ZP, PVS, and PB volumes were expressed relative to ooplasm volume. For the ZP, we additionally computed a local thickness distribution as the radial distance between the ZP outer boundary and the PVS outer boundary along matched spherical directions from the ZP geometric center. To reduce artefactual zero-thickness estimates near poorly segmented axial extremes, only directions for which both ZP and PVS boundary points lay within the central 80% of the z range were included. The mean of this thickness distribution was retained as the per-oocyte ZP thickness feature.

Global shape was characterized by elongation and spherical-harmonic descriptors. We defined elongation as 1 – *a*/*b*, with *a* the minor and *b* the major principal-axis length from 3D shape analysis (axis ordering fixed consistently throughout). Higher-order global shape features were extracted by spherical harmonic decomposition,

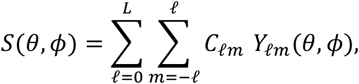

and the rotation-invariant second-degree shape power, defined as

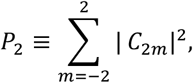

To remove size dependence, *P*_*2*_ was normalized by the zero-degree power:.

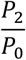

The normalized value *P*_*2*_ was used to in this analysis to quantify deviations from spherical symmetry independently of the oocyte’s orientation in the imaging frame^60^

### Dry mass and dry mass density estimation

We estimated dry-mass-related features from the reconstructed RI values, converting each voxel’s RI to a dry mass concentration through the standard linear relationship between RI increment and biomolecular concentration, *c* = (*n - n*_*m*_) / α, with *n* the voxel RI, *n*_*m*_ the RI of the surrounding medium, and α the RI increment. The conversion follows the standard quantitative phase imaging convention for converting RI contrast into biomolecular dry mass^61,62^. Dry mass density is the mean mass concentration within the compartment of interest, and total dry mass is the integral of mass concentration over the compartment volume. We used *n*_*m*_ = 1.337 (culture medium) and α *=* 0.18mL g^− 1^; all dry-mass calculations were performed using custom Python scripts. Reported dry-mass density values are reproducible within the platform but their absolute magnitudes scale linearly with the assumed values of α and *n*_*m*_; cross-platform comparisons therefore require matched calibration. The reported dry mass density values should therefore be interpreted as RI-derived mass-equivalent concentration relative to the surrounding M16 medium, rather than as absolute intracellular dry mass concentration; absolute calibration against external standards (for example, reference solutions or beads of known RI/concentration) was not performed in the present study, and inter-instrument comparisons will require such calibration^63^. However, Mean RI and dry-mass density may therefore not exhibit an exact linear correspondence in this study. For dry-mass calculations, voxel RI values below *n*_*m*_ were set to *n*_*m*_, equivalent to setting negative concentration estimates to zero, whereas mean RI was calculated directly from the reconstructed RI distribution without clipping. Importantly, the aging-associated group trend in dry-mass density was preserved when the calculation was repeated without this clipping constraint (Supplementary Fig. S8)

### RI statistical features

For the HT panel we computed three compartment-level RI-distribution descriptors (mean, SD, and CV of RI) which together constitute the RI-statistics family of the 160-feature panel (**Fig. 5**). For the modality-comparison analysis (**Fig. 7**), we additionally computed three matched voxel-intensity descriptors (mean, SD, CV of BF intensity) over the same compartment mask for the 2D BF and 2.5D BF panels; these BF intensity descriptors are used only in the modality-comparison analysis and do not appear in the HT 160-feature panel.

### Texture feature extraction from RI maps

Intracellular texture was quantified from RI maps using Haralick-style GLCM analysis. For HT data, RI values within the ooplasm mask were scaled by a factor of 10,000 and retained as integer grey levels without further quantization, so that the number of effective grey levels reflected the native RI dynamic range of each volume (typically several hundred levels per oocyte) rather than being clipped to a fixed range. For BF data, intensities were quantized to 64 grey levels following convention. GLCMs were computed volumetrically over the full 3D ooplasm mask for HT using 13 symmetry-equivalent 3D directions; for 2D BF, GLCMs were computed on the in-focus slice using four in-plane directions; and for 2.5D BF, the 2D four-direction computation was applied to each z slice and the resulting GLCMs were merged across the stack^64^. GLCMs were computed at an offset distance of 1 voxel. For each GLCM we extracted sixteen descriptors — angular second moment (ASM), contrast, correlation, variance, homogeneity, sum average, sum variance, sum entropy, entropy, difference variance, difference entropy, information measure of correlation 1 (IMC1), information measure of correlation 2 (IMC2), dissimilarity, cluster shade, and cluster tendency, yielding a compartment-level texture profile for each oocyte and imaging modality^65^.

This asymmetric quantization scheme, native integer-rounded RI levels for HT versus 64 grey levels for BF, was chosen to preserve each modality’s native dynamic range rather than to artificially equalize feature dimensionality. Imposing a 64-level cap on HT RI volumes would discard the sub-millidegree RI gradients that constitute the primary source of intracellular contrast in HT and would bias the comparison against the modality whose contrast it is intended to characterize; conversely, the 64-level BF convention reflects the histogram support typical of BF intensity images. Haralick descriptors are therefore compared as modality-native readouts at each modality’s informational ceiling. Voxel spacing was anisotropic (0.113 µm laterally and 0.549 µm axially), so a 1-voxel offset along the 13 symmetry-equivalent 3D directions mixes physical distances; texture features should accordingly be interpreted as platform-native readouts rather than physically isotropic measurements. Because several Haralick descriptors, including entropy, sum and difference entropy, contrast, and variance, increase with the number of grey levels, their absolute magnitudes are not directly comparable across the two quantization schemes; cross-modality interpretation therefore rests on which modality’s texture family tracks aging and on within-modality monotonic trends, rather than on absolute texture magnitudes.

### Population-level feature analysis

For population-level analysis, we grouped MII oocytes by aging condition (fresh, aged-24 h, aged-48 h) and split features into morphology-based and non-morphology-based groups. Morphology-based features captured compartment geometry; non-morphology-based features covered dry mass, RI statistics, and intracellular texture. Feature distributions were compared across the three aging groups to identify progressive biophysical changes associated with post-ovulatory aging. Per-group sample sizes were *n* = 96 fresh (0 h), *n* = 77 aged-24 h, and *n* = 49 aged-48 h MII oocytes.

### Comparative analysis of 2D BF, 2.5D BF, and HT-derived features

To compare the information content of different imaging modalities, we constructed three modality-specific feature sets. The 2D BF set was extracted from a single in-focus BF image; the 2.5D BF was constructed by sampling BF images from the full 3D stack at approximately 15 µm axial intervals, approximating clinically typical acquisition spacing; and the HT set was extracted from the full 3D RI volume and associated compartment masks. Low-dimensional embeddings were generated using UMAP (implemented via the MATLAB Add-On) with *n*_neighbors = 15, min_dist = 0.10, and the Euclidean metric under a fixed random seed. Group separation was quantified by the silhouette score (computed with squared Euclidean distance, MATLAB default) and by PERMANOVA *R*^2^ (computed directly on the raw feature matrix with 999 permutations). To assess the relative importance of different feature classes, we performed feature-family ablation, removing one group of features at a time and recomputing discrimination performance. The change in performance (Δ*R*^2^ – *R*^2^ _full features_ – *R*^2^ _without_) was used to estimate the contribution of morphology, intensity/RI statistics, texture, and mass-related features to modality-specific group discrimination. We note that this three-set comparison is not a strictly modality-controlled benchmark: the 2D BF, 2.5D BF, and HT feature sets differ in spatial dimensionality, feature count (28 vs 28 vs 39 ooplasm features, respectively), grey-level quantization (64-level BF vs native RI-scaled integer levels for HT), and the morphology descriptors accessible to each modality. The reported gains therefore reflect the combined effects of imaging contrast, dimensionality, and feature-engineering choices accessible to each modality, rather than RI contrast in isolation; feature-count-matched subsampling, equalized quantization sensitivity analyses, and supervised cross-validated classification were not implemented in the present study and remain natural extensions^66–68^.

### Software and code availability

Image processing, feature extraction, and statistical analyses were performed in Python and MATLAB. The AI-assisted segmentation pipeline used the pre-trained SAM 2 foundation model in its video predictor mode. Source code for segmentation prompting, preprocessing, post-processing, feature extraction, and downstream analysis is publicly available in the accompanying GitHub repository and archived on Zenodo repository, together with representative 3D RI volumes, at https://doi.org/10.5281/zenodo.20175228.

### Statistical analysis

Quantitative data are presented as mean values with individual data points overlaid, as shown in the relevant figure panels (**Fig. 6, Supplementary Figs. S2–S6**). The number of oocytes analyzed in each group is reported in the *Population-level feature analysis* section. For each of the 39 features in the HT feature set, between-group comparisons were performed using one-way ANOVA followed by Tukey’s multiple-comparison test, where appropriate. A *p*-value < 0.05 was considered significant. Exact tests, sample sizes, and significance definitions are provided in the corresponding figure legends. Feature-level pairwise comparisons across the 39 ooplasm features (Fig. 6) and the 16-feature Haralick panels (Supplementary Figs. S4–S6) are presented as exploratory descriptors of group-wise differences; per-feature P values were not corrected for the family of tested features. Oocytes from the same mouse, replicate, and culture drop were treated as independent observations in the group-wise statistics, PERMANOVA, and silhouette analyses; hierarchical mixed-effects models accounting for these nested sources of variance were not implemented in the present study. Centroid versus dispersion contributions to the PERMANOVA result were not separately tested via PERMDISP and may jointly contribute to the reported R^2^ values.

## Acknowledgements

This work was supported by National Research Foundation of Korea grant funded by the Korea government (MSIT) (RS-2024-00442348, RS-2024-00440577), Korea Institute for Advancement of Technology (KIAT) through the International Cooperative R&D program (P0028463), Commercialization Promotion Agency for R&D Outcomes (COMPA) funded by the Ministry of Science and ICT(MSIT) (RS-2024-00440577, RS-2026-25535856).

## Author contributions

J.H.K. and Y.K.P. conceived and designed the study. M.K., C.L., and J.D. performed the measurements. M.K. analyzed the data. T.S., D.L., K.H.C., S.Y.S., J.K., T.P., and C.H. contributed to data review and interpretation. All authors reviewed, edited, and approved of the final manuscript.

